# Single-section spatial hypoxia–cytotoxic associations do not consistently reproduce across breast cancer patients

**DOI:** 10.64898/2026.06.13.732045

**Authors:** Binyu Dong, Zhaoqi Song, Yangyang Yin

**Author notes:** These authors contributed equally to this work (Binyu Dong, Zhaoqi Song).

## Abstract

Spatial transcriptomics can reveal localized tumor–immune relationships, but thousands of spots from one tissue section do not provide thousands of biological replicates. We evaluated the distinction between within-section association and patient-level reproducibility using public breast cancer datasets. In a 10x Genomics Visium discovery section containing 3,798 spots, hypoxia-related transcription was inversely associated with cytotoxic gene activity in neighboring spots (Spearman *ρ* = −0.202). High-hypoxia spots also had lower neighborhood cytotoxic scores than low-hypoxia spots (rank-biserial effect = −0.286). We then tested the directional association in an independent HER2-positive cohort comprising 36 sections, 13,619 spots, and eight patients. Only 19 of 36 sections and five of eight patients showed negative associations. The median patient-level correlation was −0.043 and did not differ from zero in a one-sided exact Wilcoxon test (*P* = 0.473). Sensitivity analyses using alternative cytotoxic and hypoxia signatures, neighborhood sizes, and Kendall correlation did not support a consistent inverse patient-level effect. Thus, a strong single-section association did not consistently reproduce across patients. These results caution against interpreting spot-level spatial associations from one section as patient-level biological effects.

## 1 Introduction

Spatial transcriptomics retains tissue coordinates while measuring gene expression, enabling localized tumor–immune relationships to be studied in intact tissue [1–3]. Analytical frameworks such as SpatialDE, SPARK, Giotto, and Squidpy detect spatial expression patterns from hundreds or thousands of measurement locations [4–7]. However, the apparent sample size of a spatial-transcriptomic analysis can be misleading. Spots from one section are spatially dependent observations from one biological specimen, not independent biological replicates. Treating them as independent evidence for a patient-level claim is a form of pseudoreplication [8].

This distinction matters because spot-level significance and patient-level reproducibility answer different questions. Thousands of spots can precisely describe one section while providing little evidence about consistency across patients. A small within-section probability can therefore coexist with weak, heterogeneous, or reversed patient-level effects. External validation must preserve the patient as the biological replication unit and report discordant results rather than pool them away.

Breast cancers provide a relevant test case because they contain substantial spatial heterogeneity within and between patients. Spatial studies have localized malignant, stromal, and immune states and identified tumor-associated interactions obscured by non-spatial profiling [9–11]. Hypoxia is a plausible correlate of local immune organization because it can alter metabolism, extracellular-matrix remodeling, checkpoint signaling, and immune-cell function [12–14]. Cytotoxic lymphocytes are central effectors of antitumor immunity [15], but their abundance and transcriptional activity vary across tumor niches.

Here, we used a two-stage design to examine the spatial relationship between hypoxia-related transcription and cytotoxic gene activity. We first conducted an exploratory analysis of one public Visium breast cancer section. We then tested the directional hypoxia–cytotoxic association in the independent multi-section HER2-positive cohort reported by Andersson *et al*. [10]. The discovery section showed a clear inverse association, whereas the external cohort revealed substantial between-patient heterogeneity and no consistent patient-level effect.

## 2 Results

### 2.1 Prespecified gene programs showed spatial structure in the discovery section

After quality control, the discovery dataset contained 3,798 spots and 36,601 measured genes. We calculated six prespecified transcriptional programs: tumor epithelial, hypoxia, cytotoxic lymphocyte, myeloid, fibroblast/extracellular matrix (ECM), and epithelial–mesenchymal-transition (EMT)-regulatory programs. The EMT-regulatory set excluded collagen genes to reduce direct overlap with the fibroblast/ECM program.

All six programs showed positive spatial autocorrelation on a symmetrized six-nearest-neighbor graph. Tumor-epithelial and fibroblast/ECM programs showed the strongest organization (Moran’s *I* = 0.613 and 0.600), followed by myeloid (0.548), EMT-regulatory (0.452), cytotoxic (0.370), and hypoxia (0.278) programs. Each observed value exceeded those from 499 random permutations (empirical *P* = 0.002; Fig. 1).

**Figure 1:**
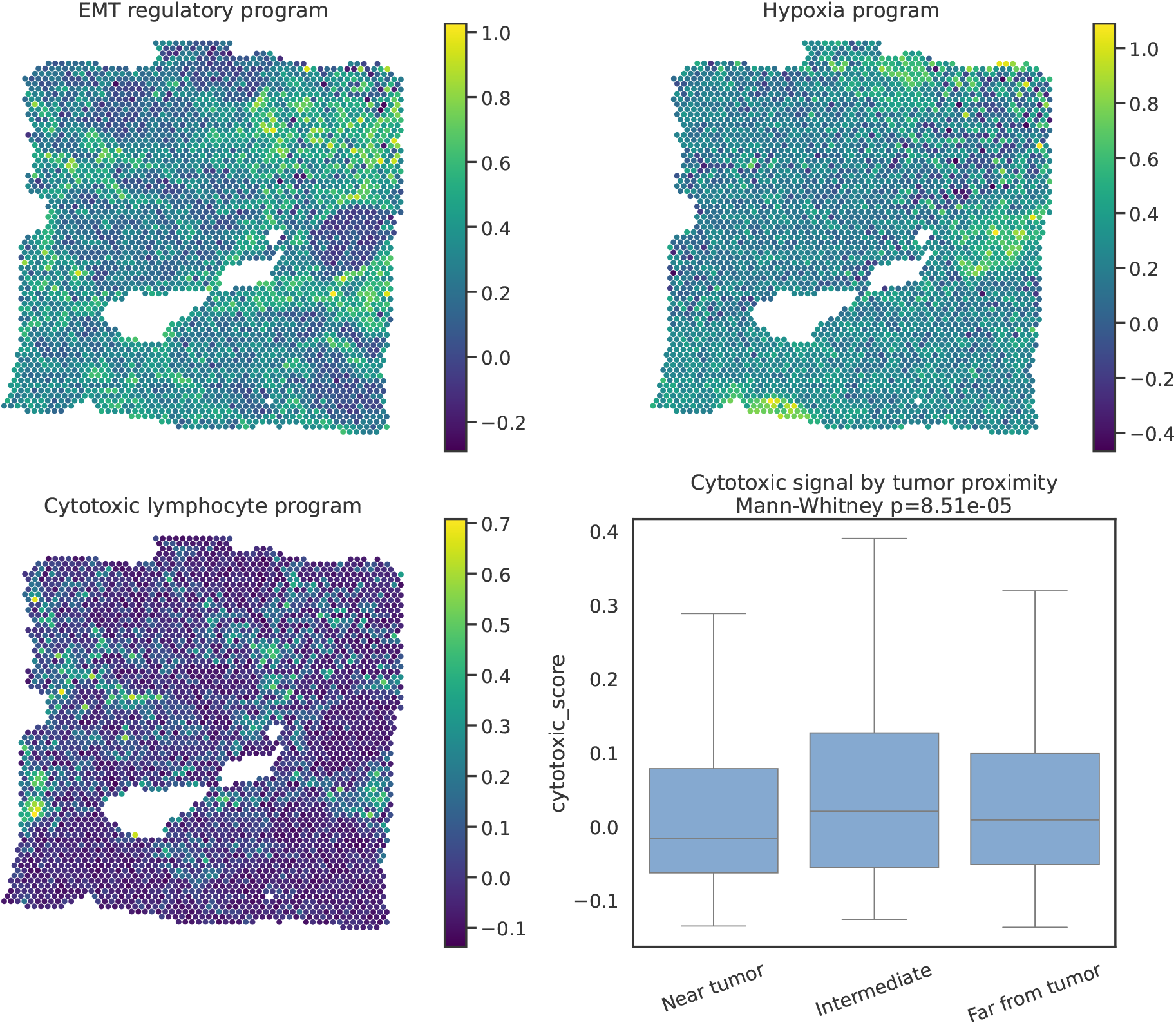
Spatial programs and tumor-proximity analysis in the discovery section. Spatial maps show EMT-regulatory, hypoxia, and cytotoxic lymphocyte program scores. The box plot compares cytotoxic scores among non-tumor spots stratified by distance to high tumor-epithelial-score spots. The displayed probability is a within-section Mann–Whitney contrast.

### 2.2 Discovery analysis identified an inverse hypoxia–neighborhood cytotoxic association

Hypoxia scores were inversely associated with the mean cytotoxic score of neighboring spots (Spearman *ρ* = *−*0.202; false-discovery-rate-adjusted *P* = 2.89 *×* 10^*−*36^). The median neighborhood cytotoxic score was −0.0147 around high-hypoxia spots and 0.0270 around low-hypoxia spots. This contrast corresponded to a rank-biserial effect of −0.286 (within-section Mann–Whitney *P* = 4.25 *×* 10^−27^; Fig. 3).

Non-tumor spots near high tumor-epithelial-score regions also had lower cytotoxic scores than distant spots. The median cytotoxic score was −0.0174 near high-score regions and 0.0080 in distant regions (rank-biserial effect = −0.102; within-section Mann–Whitney *P* = 8.51*×* 10^−5^; Fig. 1). The direction was stable when high tumor-epithelial-score regions were defined using the 70th, 75th, or 80th percentile, with rank-biserial effects from −0.102 to −0.128 (Fig. 3).

### 2.3 EMT-regulatory activity was not associated with cytotoxic exclusion

The fibroblast/ECM score was strongly associated with the neighborhood EMT-regulatory score (Spearman *ρ* = 0.730). By contrast, the EMT-regulatory score was positively associated with local cytotoxic scores (*ρ* = 0.298) and neighborhood cytotoxic scores (*ρ* = 0.340; Fig. 2). The discovery section therefore did not support the initial hypothesis that the collagen-excluded EMT-regulatory program marked cytotoxic exclusion.

**Figure 2:**
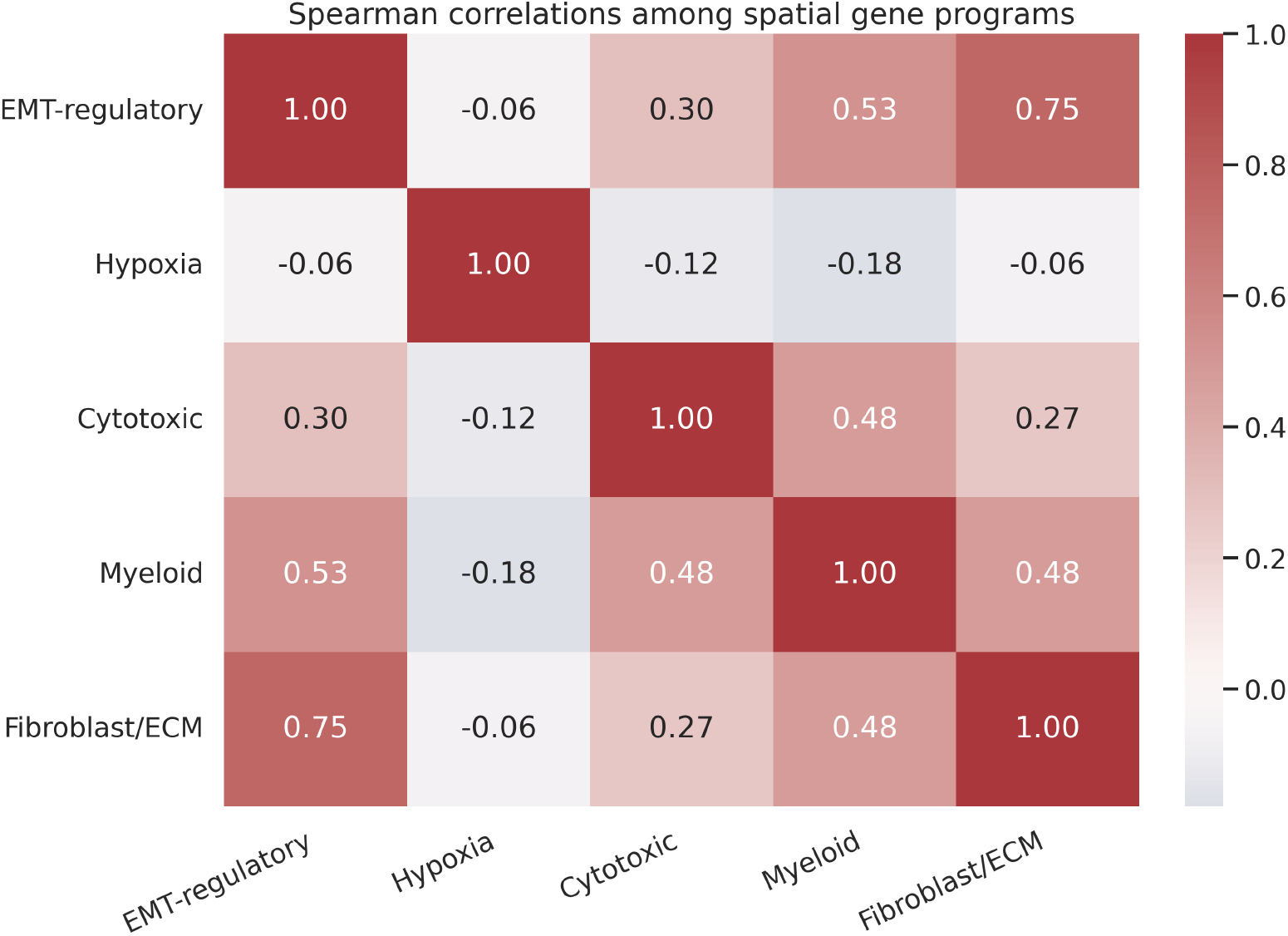
Correlations among spatial gene programs in the discovery section. The heat map shows pairwise Spearman correlations among EMT-regulatory, hypoxia, cytotoxic, myeloid, and fibroblast/ECM program scores.

**Figure 3:**
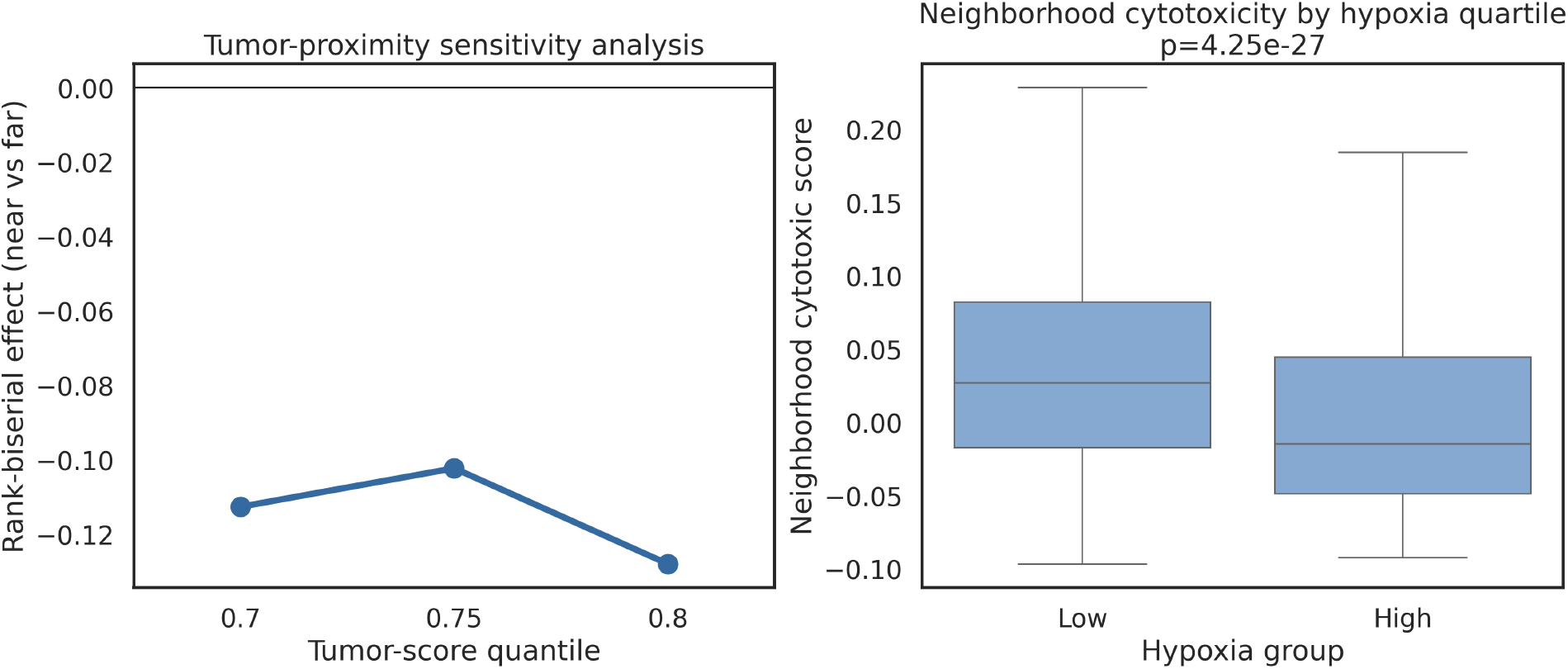
Threshold robustness and hypoxia-associated neighborhood cytotoxicity in the discovery section. Left, rank-biserial effects for near-versus-far cytotoxic contrasts using three tumor-epithelial-score thresholds. Right, neighborhood cytotoxic scores around spots in the lowest and highest hypoxia quartiles.

### 2.4 Patient-level validation did not support a consistent inverse association

We next evaluated the directional hypoxia–cytotoxic association in the Andersson HER2-positive breast cancer cohort. The external dataset contained 36 sections from eight patients and 13,619 spatial spots. Within each section, expression was library-size normalized, log-transformed, and converted to gene-wise standardized scores before calculating hypoxia and cytotoxic programs. The section-level Spearman correlation between hypoxia and neighborhood cytotoxic scores ranged from −0.317 to 0.475 (Fig. 4).

**Figure 4:**
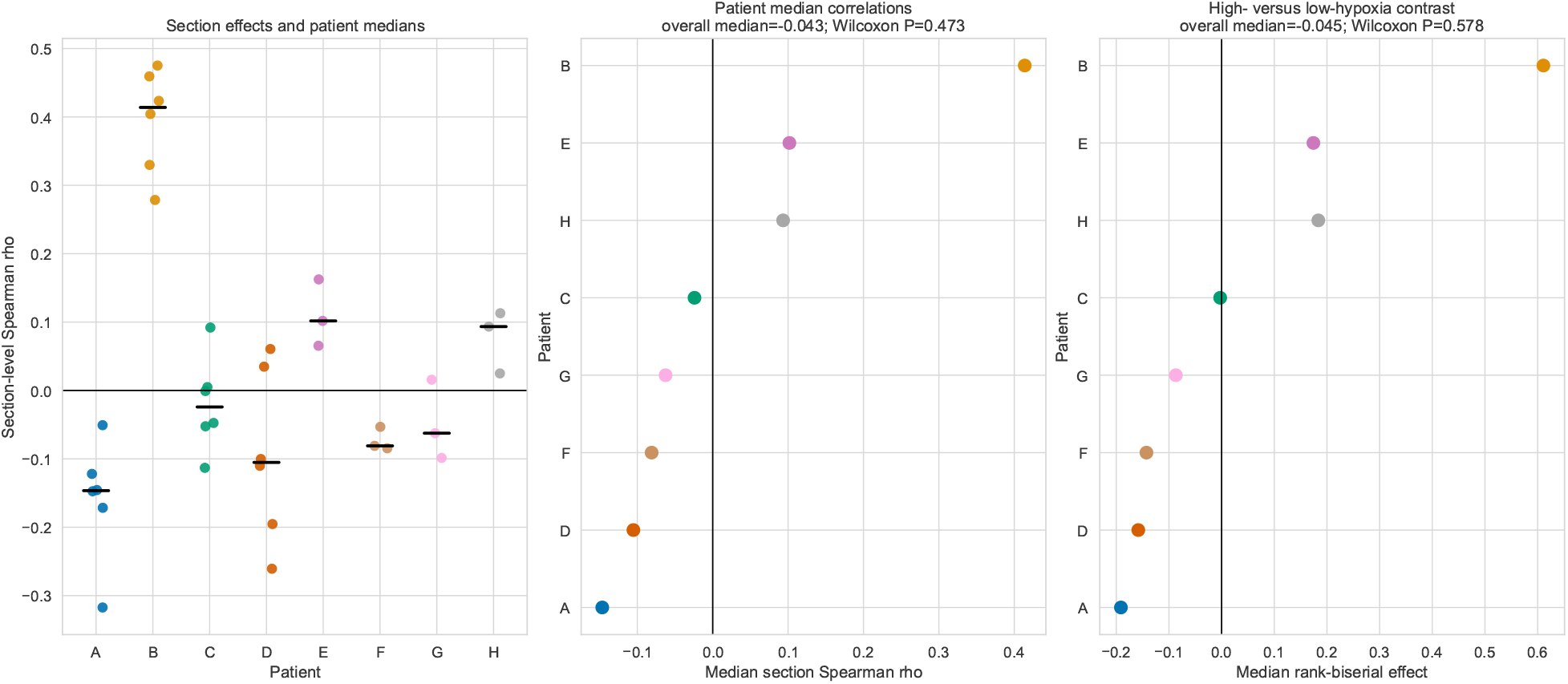
Patient-level validation does not support a consistent inverse hypoxia– cytotoxic association. Left, section-level Spearman correlations in 36 sections from eight patients; colors identify patients and black lines show patient medians. Middle, patient median section-level correlations. Right, patient median rank-biserial effects comparing high- and low-hypoxia spots. Vertical lines mark zero, and titles report the overall patient median and one-sided exact Wilcoxon probability.

Nineteen of 36 sections had negative correlations. At the patient level, five of eight patients had negative median section effects, whereas three had positive effects. The median patient-level correlation was −0.043 and was not lower than zero in a one-sided exact Wilcoxon test (*P* = 0.473). The corresponding patient-level median rank-biserial effect for high-versus low-hypoxia spots was −0.045 (*P* = 0.578). Thus, the inverse association observed in the discovery section was present in subsets of the external cohort but was not consistently reproduced across patients.

### 2.5 Sensitivity analyses reinforced the absence of a consistent inverse patient-level effect

The lack of a consistent inverse patient-level association was not explained by the six-neighbor graph or the full cytotoxic gene set. Using 4, 8, or 10 neighbors produced median patient-level Spearman correlations of −0.028, −0.025, and −0.032, respectively. Five of eight patients had negative effects under each definition, and one-sided exact Wilcoxon probabilities ranged from 0.371 to 0.473 (Fig. 5). An effector-only cytotoxic program comprising *NKG7, GNLY, GZMB*, and *PRF1* produced a median patient effect of −0.025 (*P* = 0.629). Replacing Spearman correlation with Kendall correlation produced a median patient effect of −0.033 (*P* = 0.473).

**Figure 5:**
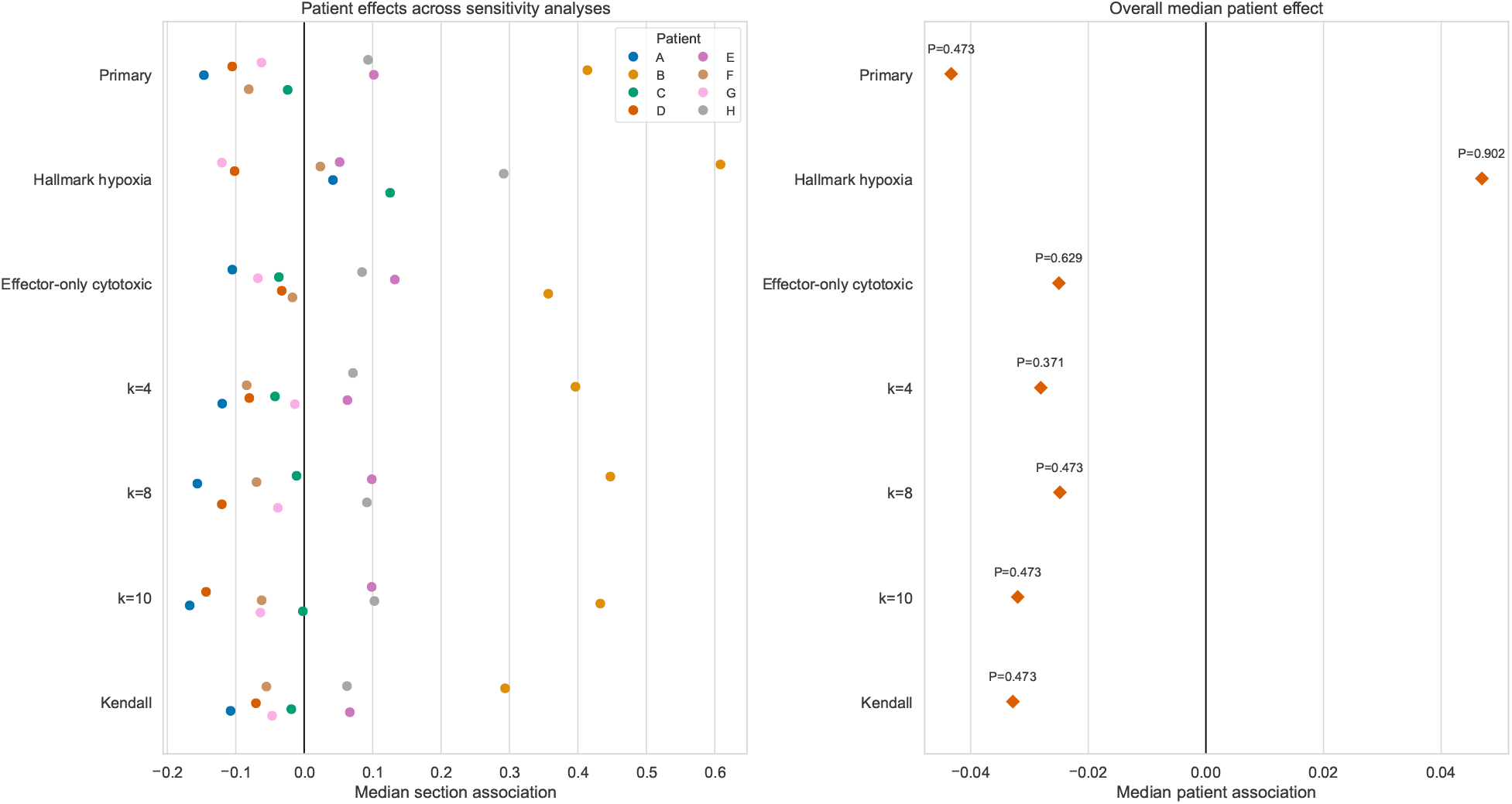
Sensitivity analyses reinforce patient heterogeneity and reveal signature dependence. Left, patient median section-level associations under the primary analysis, the MSigDB Hallmark hypoxia signature, an effector-only cytotoxic signature, alternative neighborhood sizes, and Kendall correlation. Right, overall median patient effects with one-sided exact Wilcoxon prob-abilities. No tested definition supported a consistent inverse patient-level association; the Hallmark hypoxia signature changed the median direction.

The result was more sensitive to the hypoxia signature. Replacing the compact six-gene program with the 200-gene MSigDB Hallmark hypoxia set [16] produced a positive median patient correlation of 0.047. Only two of eight patients had negative median effects under this definition (*P* = 0.902). These analyses did not identify a robust inverse association. Instead, they showed that the estimated direction depended partly on the transcriptional definition of hypoxia while remaining heterogeneous across patients.

## 3 Discussion

This study separates a strong within-section spatial association from its patient-level reproducibility. In the discovery section, hypoxia-related transcription marked neighborhoods with lower cytotoxic gene activity. However, the same directional association varied markedly across an independent multi-section cohort and was not consistently negative at the patient level. The combined evidence therefore supports spatial context dependence rather than a general hypoxia-associated cytotoxic-exclusion pattern across breast cancers. Patient heterogeneity should be treated as a biological result rather than a technical nuisance.

The discovery result remains biologically plausible. Hypoxia can reshape tumor immunity through metabolic competition, altered checkpoint signaling, and impaired immune-cell function [12–14]. Yet transcriptome-only spatial associations cannot distinguish these mechanisms from differences in cell abundance, stromal composition, tumor architecture, or section quality. The positive associations observed in several external patients show that these competing explanations are not minor technical details. They can change the direction of the measured relationship.

The external analysis also illustrates why patient-level validation should accompany spot-level inference. The discovery section produced extremely small spot-level probabilities because it contained thousands of spatial observations. Those probabilities did not predict consistency across eight independent patients. Patient B showed uniformly positive section-level associations, whereas patient A showed uniformly negative associations. This structure would be obscured by pooling all spots or treating sections from the same patient as independent replicates.

The sensitivity analyses sharpen this interpretation. Alternative neighborhood sizes, an effector-only cytotoxic program, and Kendall correlation retained weak and heterogeneous patient effects. In contrast, the Hallmark hypoxia signature changed the overall median direction. This signature dependence indicates that “hypoxia” is not a single directly measured variable in transcriptome-only analyses. The compact and Hallmark programs capture overlapping but non-identical biological processes, and neither provided evidence for a general inverse patient-level association.

### Measurement limitations

Gene-program scores are transcriptional proxies rather than direct measurements of oxygen tension, immune-cell density, or killing activity. The compact hypoxia program was selected for interpretability, whereas the Hallmark set captures a broader response program [16]. Neither was validated against direct oxygen or protein measurements in these sections.

### Cellular-resolution limitations

Visium and earlier spatial-transcriptomic spots contain mixtures of cells. Program scores can therefore combine differences in cell abundance with changes in per-cell transcriptional activity. Spatial deconvolution can reduce this ambiguity but introduces reference and model assumptions [17].

### Annotation and platform limitations

The tumor-proximity analysis used an expression-based tumor proxy rather than pathologist-defined regions. The discovery and validation datasets were also generated with different spatial-transcriptomic implementations and were normalized independently. These differences may contribute to heterogeneous effect estimates.

### Generalizability limitations

The external cohort included eight HER2-positive patients. It provides a meaningful patient-level validation step but cannot represent all breast cancer subtypes or explain treatment-associated heterogeneity. The observational design also cannot establish that hypoxia causes cytotoxic exclusion.

The most effective next upgrade is no longer simply to add more spots. Future work should add more independent patients and prespecify patient-level estimands. It should integrate histopathology, cell-type deconvolution, and spatially resolved protein measurements of hypoxia and cytotoxic effectors. A hierarchical model could estimate patient-specific effects while testing whether recep-tor status, immune composition, stromal abundance, or technical platform explains heterogeneity. These additions would convert the present reproducible association study into a mechanistic and clinically interpretable analysis.

## 4 Conclusion

A public breast cancer Visium section showed a clear inverse spatial association between hypoxia-related transcription and neighborhood cytotoxic gene activity. External testing across 36 sections from eight patients did not reproduce a consistent patient-level effect and instead revealed substantial heterogeneity and signature dependence. The study provides a reproducible example of why spot-level spatial associations should not be interpreted as patient-level biological effects without patient-level validation.

## 5 Methods

### 5.1 Study design and datasets

The study used a two-stage secondary-analysis design. The discovery analysis used the public 10x Genomics Visium dataset `V1_Breast_Cancer_Block_A_Section_1`. The filtered feature-barcode matrix file was `V1_Breast_Cancer_Block_A_Section_1_filtered_feature_bc_matrix.h5`, and spatial coordinates and images were supplied in `V1_Breast_Cancer_Block_A_Section_1_spatial. tar.gz`. Both files were downloaded from the public 10x Genomics content-delivery endpoint linked in Data availability. The external analysis used count matrices and spatial coordinates from the HER2-positive breast cancer cohort of Andersson *et al*. [10, 18]. The Zenodo archive contained one `count-matrices/<section>.tsv.gz` file and one `<section>_selection.tsv.gz` coordinate file per section. The external cohort comprised 36 sections from eight patients, identified as A–H, with 176–712 analyzed spots per section. No new human-subject recruitment or intervention was performed.

### 5.2 Discovery-section preprocessing and program scoring

The discovery data were read using Scanpy 1.11.1 [19]. Spots were retained when they contained at least 300 detected genes and less than 25% mitochondrial counts. Counts were normalized to 10,000 counts per spot and log-transformed. Gene-program scores were calculated with Scanpy’s score_genes function using random seed 20260613. The analysis used the Scanpy 1.11.1 default control-gene settings. Genes absent from the expression matrix were omitted before scoring, and the genes used were recorded in the generated results. The prespecified programs were tumor epithelial (*EPCAM, KRT8, KRT18, KRT19, MUC1*), hypoxia (*HIF1A, CA9, VEGFA, LDHA, SLC2A1, ENO1*), cytotoxic lymphocyte (*CD3D, CD3E, CD8A, CD8B, NKG7, GNLY, GZMB, PRF1*), myeloid (*LYZ, LST1, TYROBP, FCER1G, CTSS, AIF1*), fibroblast/ECM (*COL1A1, COL1A2, COL3A1, DCN, LUM, FAP, PDGFRA*), and EMT-regulatory (*VIM, FN1, SNAI1, SNAI2, TWIST1, ZEB1, ZEB2*).

### 5.3 Spatial graph and discovery-section statistics

A six-nearest-neighbor graph was constructed from spatial coordinates and symmetrized. The spatial-lag score was the mean program score among connected neighbors. Moran’s *I* quantified global spatial autocorrelation, with empirical probabilities calculated from 499 random permutations [20]. Prespecified associations were quantified using Spearman correlation, and five correlation probabilities were adjusted using the Benjamini–Hochberg procedure [21]. Upper- and lower-quartile contrasts used two-sided Mann–Whitney tests and rank-biserial effects. These probabilities describe within-section contrasts and are not patient-level inference.

### 5.4 Tumor-proximity analysis

High tumor-epithelial-score spots were defined using the upper quartile of the tumor-epithelial score. Euclidean distance from each spot to the nearest high-score spot was calculated. Remaining spots were divided into near, intermediate, and far strata using the 33rd and 67th percentiles of non-tumor distances. Sensitivity analysis repeated the procedure using the 70th, 75th, and 80th percentiles to define high-score regions.

### 5.5 External validation

Each Andersson section was analyzed separately. Spot counts were normalized to 10,000 counts and log-transformed. For each hypoxia and cytotoxic gene, expression was standardized across spots within the section. Program scores were the mean standardized expression of the prespecified genes. Section-wise processing was used to prevent sequencing-depth and section-level distribution differences from directly determining program scores. All compact hypoxia and cytotoxic genes were present in every external section. A symmetrized six-nearest-neighbor graph was constructed from the published section coordinates, and neighborhood cytotoxic scores were calculated as in the discovery analysis.

The directional validation estimand, specified before external analysis, was the Spearman correlation between compact hypoxia and neighborhood cytotoxic scores within each section. Section effects were summarized by their median within each patient. The patient median was selected as the primary biological estimand because sections from the same patient are repeated observations and because the median limits the influence of one atypical section. The primary patient-level test was a one-sided exact Wilcoxon signed-rank test of whether patient median correlations were below zero. A one-sided exact sign test quantified directional consistency. High-versus low-hypoxia quartile contrasts were evaluated as a secondary patient-level analysis.

### 5.6 External-validation sensitivity analyses

Sensitivity analyses replaced the compact hypoxia program with the 200-gene MSigDB Hallmark hypoxia set, of which 179–190 genes were present per section [16]. A second analysis replaced the full cytotoxic program with an effector-only set comprising *NKG7, GNLY, GZMB*, and *PRF1*. Additional analyses used 4, 8, or 10 nearest neighbors instead of 6 and replaced Spearman correlation with Kendall correlation. Each sensitivity analysis retained section-wise processing and patient-median aggregation. These analyses were added after the primary analysis to evaluate dependence on scoring and neighborhood definitions.

### 5.7 Reproducibility

Random seeds were fixed at 20260613 where stochastic procedures were used. The complete discovery and validation scripts, requirements, result tables, and figures accompany this manuscript.

## Data availability

The discovery dataset is available from 10x Genomics as a filtered feature-barcode matrix and a spatial archive. The external HER2-positive cohort is available from Zenodo under DOI 10.5281/zenodo.4751624.

## Code availability

Executable analysis scripts, generated result tables, and figures are publicly available at https://github.com/eughu/spatial-hypoxia-cytotoxic-validation.

## Author contributions

B.D.: Conceptualization, Data curation, and Formal analysis. Z.S.: Investigation, Methodology, Project administration, Resources, Software, and Supervision. Y.Y.: Validation, Visualization, Writing – original draft, and Writing – review and editing.

## Ethics statement

This study was a secondary analysis of publicly available, de-identified data. No new human participants were recruited, no intervention was performed, and no animal experiments were conducted. This study was not a clinical trial.

